# Characterization of the molecular mechanisms of early sexual maturation stages in the Australian greenlip abalone (*Haliotis laevigata*)

**DOI:** 10.1101/2025.08.18.670794

**Authors:** Ya Zhang, Carmel McDougall, Ido Bar, Natasha Botwright

## Abstract

Sexual maturation is a critical developmental transition in abalone, yet its molecular basis remains unclear. Understanding the mechanisms underlying maturation is essential for improving the yield and quality of aquaculture production. This study employed transcriptomic analysis to identify key genes involved in the early stages of sexual maturation in the greenlip abalone (*Haliotis laevigata*), a commercially and ecologically important species native to Australia. Among the most significantly differentially expressed genes were members of the 5-hydroxytryptamine receptor, cytochrome P450, and HSP70 families, which are associated with neuroendocrine signalling, steroid metabolism, and cellular stress response, respectively. Gene regulatory network analysis further predicted their upstream and downstream interactions, suggesting potential signalling pathways that may coordinate reproductive initiation. These findings provide new insights into the molecular mechanisms underlying early maturation in abalone and establish a foundation for future functional investigations.

## Introduction

Abalone are economically valuable marine species widely cultured for their edible and highly prized muscular foot (Cook, 2016). Abalone are also extremely important to local fishing communities (Prince, 2003). In recent years, wild abalone catch has been declining (Gordon and Cook, 2001, Cook, 2023b) due to a combination of factors. These include overexploitation, recruitment failure due to stock depletion, inadequate spatial management, diseases, increased predation, climate change, and habitat degradation (Cook, 2023a). Amongst these, illegal harvesting is likely the most significant factor driving the global fisheries decline (Cook, 2023b). Of the few remaining wild abalone fisheries, primarily located in New Zealand, Australia, Mexico, and Japan, all have experienced significant stock depletion (Cook, 2023b). To maintain supply of abalone despite the depletion of wild stock, farmed abalone production has grown rapidly since around 2000, particularly in China and South Korea (Sim et al., 2021). China is now the largest producer of abalone, with an annual production of 163,000 metric tons in 2021/22, followed by South Korea at over 20,000 metric tons (Cook, 2023b).

Expanding abalone aquaculture not only boosts production but also alleviates pressure on wild stocks by offering opportunities to restore or enhance natural populations through hatchery-produced juveniles (Cook, 2023a). The greenlip abalone (*H. laevigata*) is a temperate species mainly distributed in southern Australia (Botwright et al., 2019) and is one of the primary cultivated Australian abalone species (Dang et al., 2011).

Despite significant growth, abalone aquaculture continues to face numerous challenges (Botwright et al., 2021), particularly reliable reproduction at scale. Traditional hatchery practices rely on various methods to artificially induce spawning in abalone, including desiccation (Zhang et al., 2004, Park and Kim, 2013), thermal shock (Setyono, 2006, Henry, 1996), pH adjustment (Henry, 1996, Peña, 1986), and the use of ultraviolet irradiated seawater (Guo et al., 1993). However, these artificial spawning induction methods often yield inefficient results, with only about 40% of broodstock spawning successfully, and outcomes varying between species (Botwright et al., 2021, Kim et al., 2018, Elliott, 2000, Al-Rashdi and Iwao, 2008).

While spawning is critical for breeding efficiency, the process of sexual maturation can adversely affect cultured abalone production. During sexual maturation, abalone redirect their energy to gonad development, leading to decreased growth rates and diminished meat quality (Piferrer et al., 2009, Desrosiers et al., 1993). High temperatures during the breeding season further compound the negative effects of sexual maturation by altering water quality parameters, such as pH, dissolved oxygen, and ammonia levels (Vandepeer, 2006). Exposure to these extreme conditions frequently triggers spontaneous spawning which is energetically taxing. If abalone spawn multiple times during the peak of summer their energy reserves become depleted, their immune system may be compromised, and they become more susceptible to pathogens. This may lead to mass mortality of abalone in production (Shiel et al., 2017). In addition, stress experienced during transportation and storage of live abalone may induce gamete release, further increasing mortality rates during transit (Hermesch and Dominik, 2021).

Understanding the mechanisms of abalone gonad maturation is therefore essential for optimising breeding efficiency, reducing transport-related stress and mortality, enhancing resilience, and improving growth and meat quality. Abalone sexual maturation is influenced by both external environmental factors (Uki and Kikuchi, 1984), such as water temperature (Grubert and Ritar, 2005, García-Esquivel et al., 2007), photoperiod (García-Esquivel et al., 2007), and food availability (Wells and Keesing, 1989), as well as internal neuroendocrine mechanisms. Several neuropeptides produced by the ganglia have been shown to regulate gonadal development. For example, gonadotropin-releasing hormone (GnRH), 5-hydroxytryptamine (5-HT), and heat shock protein 70 (HSP70) have been widely reported to be associated with reproductive maturation in abalone (Nuurai et al., 2016, Kim et al., 2019a, Mendoza-Porras et al., 2017). In addition, egg-laying hormone (ELH), a neuropeptide secreted by the neural ganglia, triggers egg-laying and related behaviours. ELH has been identified in greenlip abalone (*Haliotis laevigata*) and was able to induce 80% spawning efficiency via injection under experimental conditions (Botwright et al., 2019). In Pacific abalone (*H. discus hannai*), another neuropeptide, APGWamide, has been shown to enhance spermiation and oocyte maturation (Kim et al., 2018, Chansela et al., 2008). Additionally, neuropeptides such as achatin, FMRFamide, crustacean cardioactive peptide, and pedal peptides A and B are significantly upregulated during maturation (Kim et al., 2019b). In greenlip abalone (*H. laevigata*), pedal peptides also showed early upregulation, suggesting a role in initial reproductive development (Botwright et al., 2019). Other neuropeptides, such as myomodulin and insulin-related peptide, have been shown to be upregulated during sexual maturation in abalone (*H. asinina*) (York et al., 2012), while buccalin has been reported to increase in expression in oysters (*S. glomerata*) (Van In et al., 2016). These findings suggest that sexual maturation in abalone is closely linked to ganglionic neuropeptide activity.

Beyond neuropeptides, a number of other genes have been implicated in gonad maturation in abalone. These include fatty acid-binding protein (Kim et al., 2020), alpha-amylase (Kim et al., 2020), cytochrome P450 family (Su et al., 2024), and vasa-like (Luo et al., 2024). However, key questions remain unanswered. In particular, the molecular mechanisms that initiate sexual maturation in both sexes are still not fully understood, including the specific genes and signalling pathways involved. Identifying the key regulators that induce gonad maturation will provide important insights for developing strategies to control or delay reproductive maturation in aquaculture systems.

(Botwright et al., 2019)(Dang et al., 2011)To investigate early gonad maturation, we conducted transcriptomic analyses on gonad and ganglia tissues collected from both male and female greenlip abalone at various stages of sexual maturity, assessed using the visual gonad index (VGI) (Kikuchi, 1974, Fukazawa et al., 2007). High-quality transcriptomes were generated to identify differentially expressed genes associated with early gonad maturation. By analysing gene expression across developmental stages and inferring gene regulatory networks, this study aimed to uncover key molecular pathways involved in the initiation of sexual maturation in *H. laevigata*.

## Materials and Methods

### Sampling

Samples for this study were collected from a single abalone cohort beginning pre-sexual maturation with a further 50 animals sampled every two months from 2011 to 2012 from a commercial abalone farm in South Australia, Australia (Mendoza-Porras et al., 2014). The abalone were randomly sampled at different time points during their natural sexual maturation cycle, resulting in a time-dependent progression of gonadal development stages, as determined by the visual gonad index (VGI). Due to the continuous spawning and gonad regeneration cycle during the spawning season no female abalone with late-stage VGI values (>2) were present within the samples. Ganglia and gonad tissues were dissected, snap-frozen in liquid nitrogen, and stored at -80°C for downstream processing. Table 1 provides the sample design for the current study, including the VGI and sex. Photographs of the gonads at different developmental stages are provided in Supplementary Figure S1.

**Table 1.** Distribution of gonad and ganglia tissue samples across visual gonad index (VGI) stages and sex in Haliotis laevigata.

| VGI | Sex | Number of Samples |  |
| --- | --- | --- | --- |
|  |  | Gonad | Ganglia |
| 0 | Unknown | 10 | 10 |
| 1 | Male | 5 | 5 |
| 1 | Female | 5 | 5 |
| 2 | Male | 5 | 5 |
| 2 | Female | 5 | 5 |
| 3 | Male | 5 | 5 |

Approximately 100 mg of the tissue sample was weighed into a 2 mL screw-capped tube kept on dry ice before the addition of 1 mL of TRIzol™ Reagent (ThermoFisher). The samples were homogenized with a Precellys 24 Lysis Homogenizer at 6800 rpm for 30 s, before RNA extraction following the TRIzol™ manufacturer’s instructions (Gottshall et al., 2008). The RNA pellets were air dried for 5-10 minutes and resuspended in 44 μL RNAse-free water. To remove contaminating DNA, the RNA preparations were treated with Turbo DNA Free (Ambion), and RNA clean-up was performed using the RNeasy Mini Kit (QIAGEN). The total RNA concentration was measured using a Qubit® fluorometer, and RNA integrity was assessed by agarose gel electrophoresis.

### Reference transcriptome

The current draft genome and transcriptome assembly for the greenlip abalone is based exclusively on a single female individual (Botwright et al., 2019). To achieve a comprehensive transcriptome study of sexual maturation in both male and female greenlip abalone, total RNA was pooled into two groups: gonad and ganglia (Table 1). In each pool three samples were included from the male and female individuals for each sexual maturation stage. Each pool contained 1 μg total RNA in equal concentrations. Libraries were constructed separately for each pool by Macrogen Oceania, Korea, using the TruSeq Standard mRNA Kit (Illumina) followed by 150 bp paired-end sequencing on the Novaseq 6000 platform.

Quality assessment of the raw sequencing reads was performed using FastQC (v0.11.9) (Andrews, 2025) to evaluate metrics such as base quality, GC content, and adapter contamination. Low-quality base pairs and adapter sequences were trimmed using Trim Galore (v0.6.6) (Krueger, 2021) with default settings resulting in a dataset of high-quality reads for downstream analysis. A single *de novo* transcriptome assembly was completed from gonad and ganglia reads using Trinity (v2.15.1) (Haas et al., 2013) producing a comprehensive set of assembled transcripts. These transcripts were annotated using DIAMOND (v2.0.15) (Buchfink et al., 2015) by aligning against the NCBI non-redundant (nr) database, applying an e-value cutoff of 1 x 10 . Transcriptome completeness was evaluated using BUSCO (v5.2.2) (Simão et al., 2015) with the Metazoa odb10 lineage dataset, providing a measure of assembly quality based on conserved single-copy orthologs. Raw sequences are publicly available under NCBI BioProject PRJNA1242192. Datasets for the assembled transcripts and annotation are available for download via the CSIRO Data Access Portal (Zhang et al., 2025).

### Sequencing for differential gene expression during maturation

Individual sequencing of abalone gonad and ganglia transcriptomes was performed using the CEL-Seq2 protocol, a highly sensitive single-cell transcriptomics method designed to analyze gene expression in single-cell or low-cell-count samples (Hashimshony et al., 2016). The method can also be used for cost-effective sequencing of a large number of bulk RNA samples (McDougall et al., 2021). The initial RNA/primer/ERCC/dNTP mix was prepared using 25 ng of RNA and 0.5 μL of the ERCC spike-in mix (1:10,000 dilution). Library quality was assessed using the Agilent TapeStation with High Sensitivity D1000 ScreenTape. Paired-end sequencing was conducted at the Ramaciotti Centre for Genomics, University of New South Wales, Australia. Custom sequencing was performed on a single lane of a NovaSeq 6000, with a 26 bp read 1, a 95 bp read 2, and a 6 bp index read.

### Data analysis

The CEL-Seq2 pipeline (Hashimshony et al., 2016, McDougall et al., 2021), modified to accommodate a different read length and mapping to a transcriptome, was used to generate the transcript abundance counts. These were then converted to UMI counts using the method outlined in previous studies (Hashimshony et al., 2016, McDougall et al., 2021). Gene names and isoform numbers were then separated in the initial data column using the stringr package (Wickham, 2019) in R (R Core Team, 2013) to create grouping variables for subsequent merging steps. Next, the collapseRows function from the WGCNA package (Langfelder and Horvath, 2008) was applied to consolidate multiple isoforms of each gene into a single expression value, ensuring each gene occupied a single row in the dataset.

Differential gene expression analysis was performed with the DESeq2 (v1.16.1) package (Love et al., 2014) in R. Raw read counts were imported, rounded to integers, and converted into a matrix. Sample metadata, including experimental conditions (e.g., treatment and time points), were annotated, and the reference group was set as “VGI0”. Principal component analysis (PCA) was performed on normalized, rlog-transformed expression data using the DESeq2 package, and visualized with ggplot2 (Villanueva and Chen, 2019). Differentially expressed genes (DEGs) were identified by fitting a generalized linear model to normalized counts (∼ 0 + condition), with separate comparisons conducted between VGI0 and VGI2 in females, and VGI0 and VGI3 in males. A log2FoldChange threshold of ±1 and adjusted p-value (padj) < 0.05 were used to determine differential expression. Heatmaps of the top 50 DEGs, selected based on statistical significance (padj), were generated using variance stabilizing transformation (VST)-normalized counts. Data were scaled to z-scores (Anders and Huber, 2010) to enhance visualization, and heatmaps were plotted using the pheatmap package (Kolde and Kolde, 2015). The code used to generate data and figures is available for download from the CSIRO Data Access Portal (Zhang et al., 2025).

To complement the differential expression analysis, a targeted set of candidate genes was selected based on prior studies in molluscs. These genes were chosen for their reported roles in reproductive development, neuroendocrine signalling, and gametogenesis. The species in which these genes were originally reported, along with the tissue types in which their functions have been implicated, are summarized in Supplementary Table S1. This candidate gene approach was included to ensure that biologically relevant regulators were assessed, particularly those with low expression levels or tightly regulated activity such as signalling molecules and transcription factors. These candidates were used to generate heatmaps of expression profiles across developmental stages in gonad and ganglia tissues, allowing visual comparison of their transcriptional activity in relation to sexual maturation. Heatmaps of candidate genes associated with sexual maturation were constructed using VST-normalized expression values, followed by Z-score transformation to emphasize relative expression patterns across samples.

Gene regulatory network (GRN) predictions were generated using the GRNBoost2 algorithm (Moerman et al., 2019). based on the expression profiles of significantly differentially expressed genes identified in each sex and tissue group. All analyses were performed using default parameters. For network visualization, predicted gene–gene interactions (edges) were ranked by importance score and imported into Cytoscape v3.10.2 (Shannon et al., 2003). Supplementary Files S1–S4 provide the predicted GRN edge lists for each tissue and sex combination: S1 – Female gonad, S2 – Female ganglia, S3 – Male gonad, and S4 – Male ganglia. Each file contains the full set of GRN edges derived from differentially expressed genes within the respective group. In subnetwork analyses, genes previously identified as associated with sexual maturation were selected as seed nodes, and their direct connections were examined to explore potential regulatory pathways relevant to reproductive development.

## Results

### Reference transcriptome

The greenlip abalone gonad and ganglia reference transcriptome for this study was assembled from 118,365,241 high-quality raw reads, resulting in 68,988 genes and 136,810 transcripts with a total assembled length of 164,924,718 bases. The average transcript length was 1206 bp, with a contig N50 of 1574 bp, reflecting the quality of the assembly. Of the 136,810 assembled transcripts, 43.1% were successfully annotated against the NCBI non-redundant (nr) protein database. The transcriptome assembly showed 88.3% completeness based on the BUSCO analysis using the Metazoa odb10 lineage dataset. This included 842 complete orthologs, comprising 372 single-copy and 470 duplicated BUSCOs, as well as 58 fragmented and 54 missing orthologs out of 954 expected genes, indicating a robust and comprehensive reference transcriptome.

### Differential gene expression

Gene expression in gonad and ganglia from each individual was assessed using CEL-Seq2, with an average sequencing depth of 6.66 million reads per sample (ranging from 709,867 to 17,604). In female gonads, 1,231 genes were found to be significantly differentially expressed between relatively mature (VGI2) and immature gonads (VGI0), among which 516 were upregulated in VGI2, and 715 were downregulated. The complete list of differentially expressed genes is available in Supplementary Table S2. In female ganglia, a total of 19 genes were identified as significantly differentially expressed between the relatively mature and immature stages. The complete list of differentially expressed genes is available in Supplementary Table S3.

In male gonad, a total of 3,384 genes were found to be significantly differentially expressed between relatively mature gonads (VGI3) and immature gonads (VGI0), among which 2,174 genes were upregulated in relatively mature gonads (VGI3) and 1,210 genes were downregulated. The complete list differentially expressed genes is available in Supplementary Table S4. In male ganglia, a total of 165 genes were significantly differentially expressed between the relatively mature (VGI3) and sexually immature (VGI0) stages, with 164 genes upregulated in relatively mature gonads (VGI3) and only one downregulated. The complete list differentially expressed genes is available in Supplementary Table S5.

### Expression patterns of candidate maturation-related genes during female maturation

Gonad development in the female samples in this study is graded on a scale from VGI 0 to 2, with VGI 0 indicating no initiation of gonad sexual maturation, and VGI 2 more advanced sexual maturation. To investigate the transcriptional dynamics of key maturation genes during ovarian development, a heatmap was generated based on Z-score normalized expression values of candidate genes with known roles in female maturation (Figure 1). Overall, distinct stage-specific expression patterns were observed for these maturation-related genes. In female gonad, genes related to meiosis (meiosis expressed gene 1 protein homolog, meiosis-specific coiled-coil domain-containing protein MEIOC-like) and the 5-HT receptor 1B-like (5-HTR1B-like) transcript showed significantly increased expression as sexual maturation progressed. In contrast, several metabolism-related genes, including fatty acid-binding protein brain-/liver-like, alpha-amylase-like, and yolk ferritin-like, exhibited decreased expression during maturation. A number of genes associated with sexual maturation were consistently expressed across developmental stages and did not show differential expression; these include FMRFamide-like neuropeptide 7, APGWamide-like neuropeptide, transcription factor SoxB1, vasa-like, 5-HT receptor 1A-like, vitelline envelope sperm lysin receptor-like, several isoforms of cytochrome P450 3A, HSP 70 4-like, and 17-beta-hydroxysteroid dehydrogenase 14-like isoform X1. Other genes previously implicated in sexual maturation had low or undetectable expression during the early stages; these include buccalin-like, doublesex- and mab-3-related transcription factor 1-like, egg laying hormone, estradiol 17-beta-dehydrogenase 8-like, HSP 70 12A-like isoform X4, and 3 beta-hydroxysteroid dehydrogenase type 7-like.

**Figure 1.**
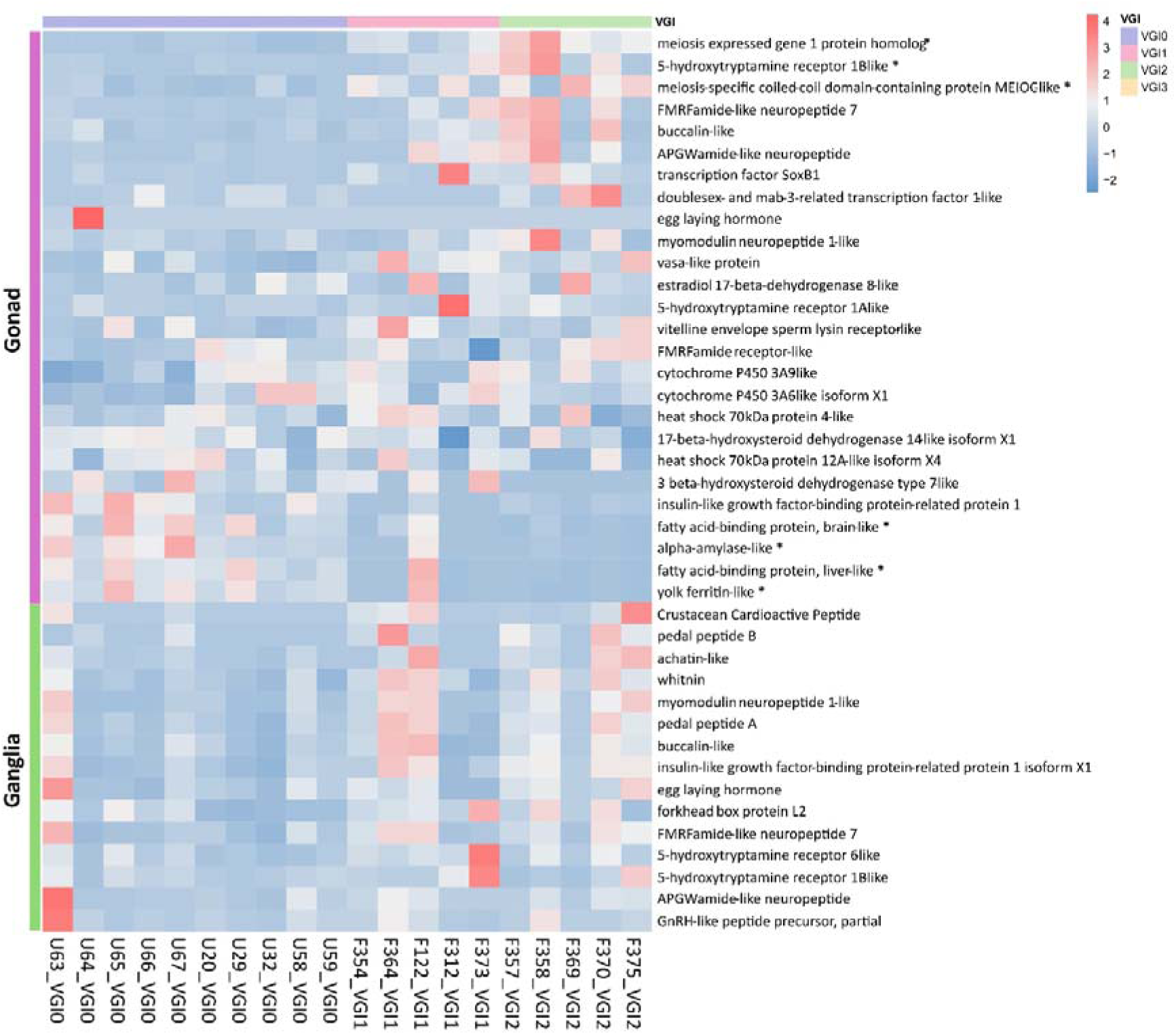
Expression patterns of maturation-related genes in female gonad and ganglia tissue across developmental stages (VGI0 to VGI2). Expression levels are color-coded, with blue indicating low expression and red indicating high expression. The developmental stages are labelled at the top of the heatmap and color-coded (VGI0: blue, VGI1: pink, VGI2: green). Asterisks indicate genes that were found to be significantly differentially expressed in the DESeq2 analysis.

In female ganglia, none of the genes of interest listed in the heatmap showed significant changes in expression across the stages of sexual maturation. However, it is noteworthy that FMRFamide-like neuropeptide 7, APGWamide-like neuropeptide, and GnRH-like peptide precursor exhibited relatively high expression levels compared to other genes and remained consistently expressed throughout the maturation process. In contrast, genes such as crustacean cardioactive peptide, pedal peptide B, whitnin, egg laying hormone, and forkhead box protein L2 showed extremely low expression levels, suggesting that they may not be involved in early-stage reproductive regulation in the ganglia. Supplementary Table S6 contains the corresponding expression data for these genes.

### Differentially expressed genes during female gonad development

The top 50 differentially expressed genes (by adjusted p-values) were illustrated in the heatmap (Figure 2). Gene expression profiles distinctly clustered between immature gonads (VGI0) and relatively mature ovaries (VGI1 and VGI2), with the exception of one VGI1 sample. The majority of these genes showed low expression levels at the immature stage but were significantly upregulated at VGI1 and VGI2, indicating a coordinated activation of transcriptional programs during ovarian maturation. Annotation of these top differentially expressed genes revealed their involvement in key biological processes, including cell cycle regulation (cell division control protein 6 homolog, kinesin-like protein KIF22, dual specificity protein kinase Ttk-like, targeting protein for Xklp2 homolog), ciliogenesis and motility (cilia- and flagella-associated protein 99-like, radial spoke head protein 9 homolog, dynein axonemal light chain 1-like, dynein axonemal light chain 4-like, centromere protein F-like), cytoskeletal organization (MORN repeat-containing protein 1-like isoform X2, armadillo repeat-containing protein 3-like), transcriptional regulation (BTB/POZ domain-containing protein 16-like isoform X2, PHD finger protein 23B-like isoform X1, LIM/homeobox protein Awh-like), transmembrane transport and metabolism (ABC transporter F family member 4-like, carbonic anhydrase 2-like, nucleoside diphosphate kinase 7-like isoform X2), and protein signaling and structural maintenance (parkin coregulated gene protein homolog, cornifelin homolog, leucine-rich repeat-containing protein 51-like). These findings reflect the tightly coordinated molecular events that drive ovarian maturation in female *H. laevigata*.

**Figure 2.**
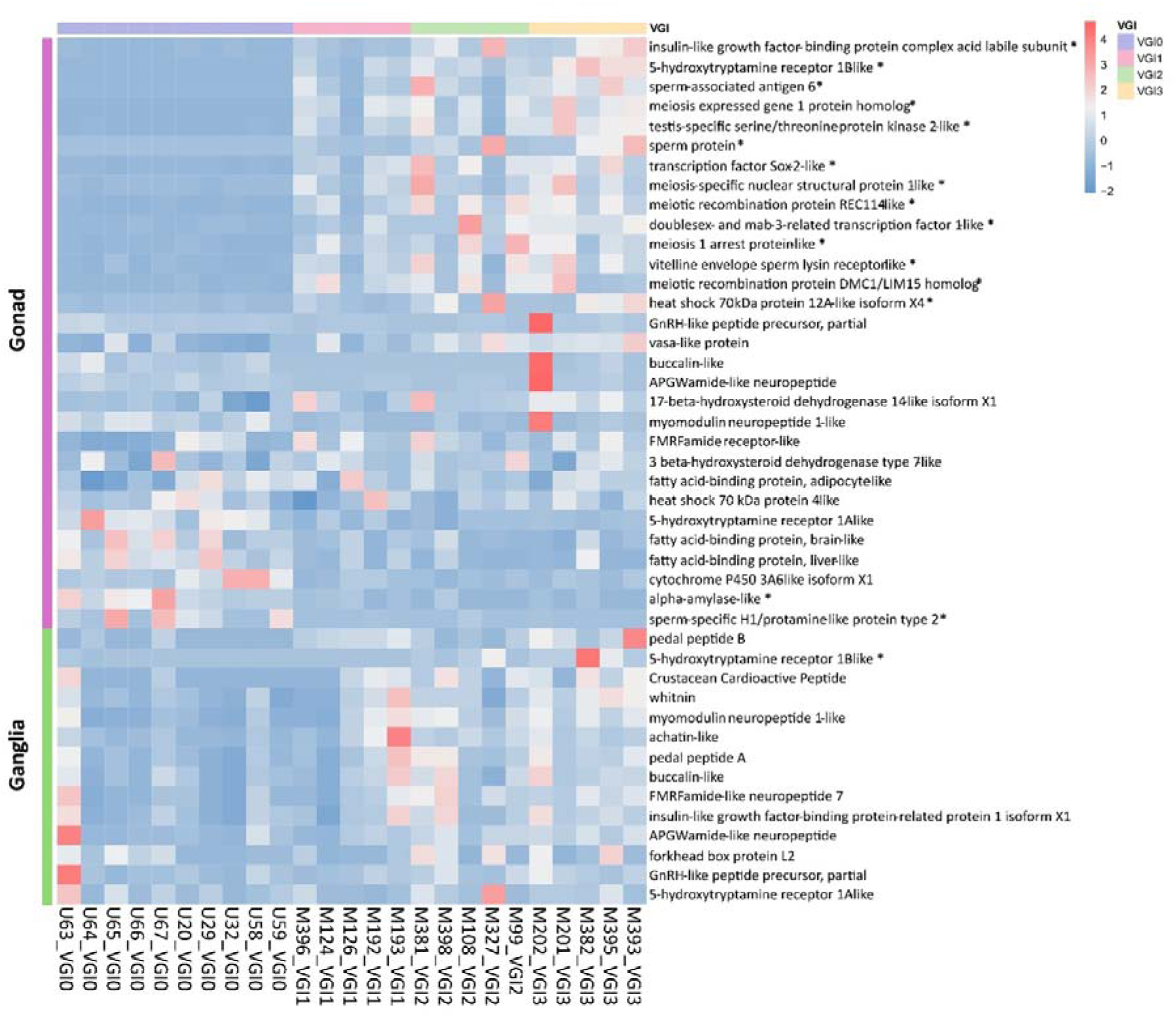
Heatmap showing the expression profiles of the top 50 most significantly differentially expressed genes (adjusted p-value) in female gonads across three developmental stages (VGI0 to VGI2). Samples are hierarchically clustered based on expression similarity. Expression levels are color-coded, with blue indicating low expression and red indicating high expression. The developmental stages are labelled at the top of the heatmap and color-coded (VGI0: blue, VGI1: pink, VGI2: green). Most genes show increased expression at VGI1 and/or VGI2 relative to VGI0, reflecting transcriptional activation during gonadal maturation.

### Differentially expressed genes in ganglia during female gonad maturation

None of the known maturation genes expressed in the female ganglia were found to be differentially expressed between VGI0 and VGI2 (Figure 1). However, 17 other genes were upregulated and 2 were downregulated during maturation, in distinct stage-specific transcript patterns (Figure 3). In addition to the sustained upregulation of cytochrome P450 3A24-like and HSP 70 B2-like during the VGI1 and VGI2 stages, other significantly upregulated genes included cell division control protein 42 homolog, BAG family molecular chaperone regulator 3-like, transmembrane protein 205-like, and DNA damage-inducible transcript 4-like protein, as well as several unannotated genes. The expression patterns of these genes suggest potential involvement in metabolic regulation, cellular stress responses, and signal transduction processes within the ganglia during reproductive maturation.

**Figure 3.**
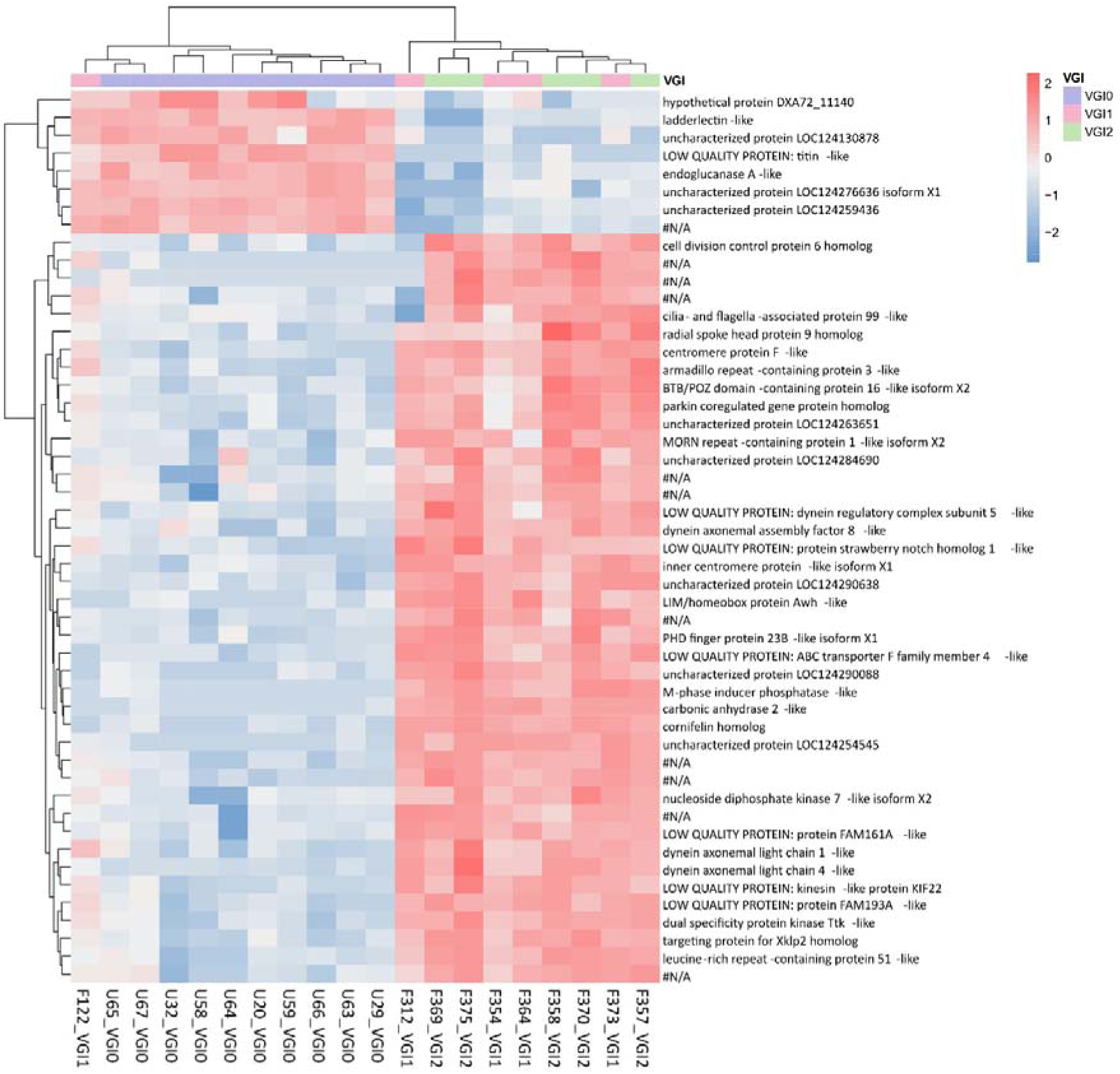
Heatmap showing the expression profiles of 19 significantly differentially expressed genes (adjusted p-value) in female ganglia between immature (VGI0) and relatively mature (VGI2) stages. Samples are hierarchically clustered based on transcript expression similarity. Expression levels are color-coded, with blue indicating low expression and red indicating high expression. Developmental stages are color-coded at the top (VGI0: blue, VGI1: pink, VGI2: green).

### Expression patterns of candidate maturation related genes during male maturation

Development in male gonads is graded on a subjective scale from VGI 0 to 3, with VGI 0 indicating no initiation of gonad sexual maturation and VGI 3 representing advanced sexual maturation. To investigate the transcriptional dynamics of key maturation genes during testis development, a heatmap was generated based on Z-score normalized expression values of genes with known roles in male maturation (Figure 4). As observed in female gonad, 5-HT receptor 1B-like (5-HT1B) was upregulated in male gonads at later maturation stages. Insulin-like growth factor-binding protein complex acid labile subunit, and sperm-associated antigen 6 were also significantly upregulated.

**Figure 4.**
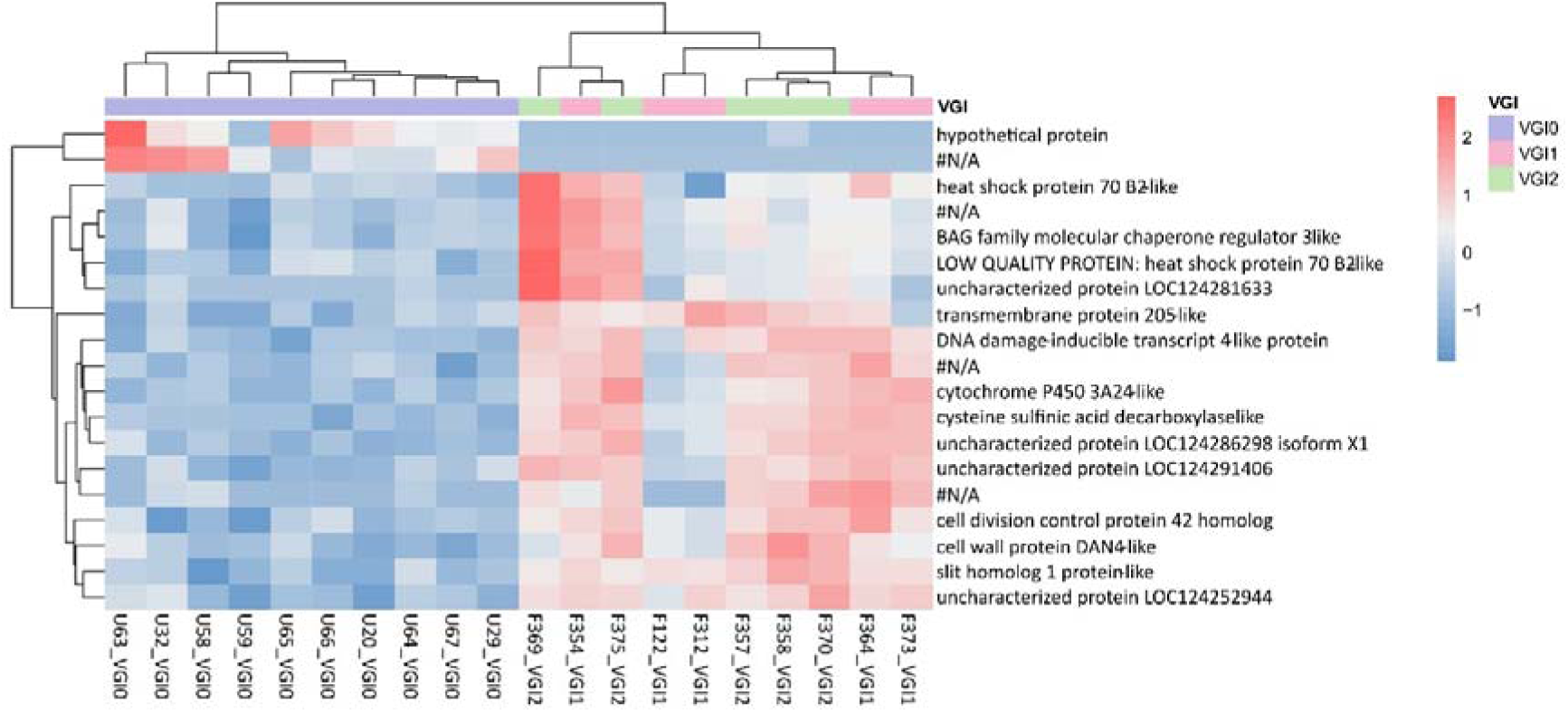
Expression patterns of maturation-related genes in male gonad and ganglion tissues across developmental stages (VGI0 to VGI3). Expression levels are color-coded, with blue indicating low expression and red indicating high expression. The developmental stages are labelled at the top of the heatmap and color-coded (VGI0: purple, VGI1: pink, VGI2: green, VGI3: orange). Asterisks indicate genes that were found to be significantly differentially expressed in the DESeq2 analysis.

Multiple meiosis-related genes, such as meiosis expressed gene 1 protein homolog, meiotic recombination protein REC114-like, and meiotic recombination protein DMC1-like, along with key transcription factors including testis-specific serine/threonine-protein kinase 2-like, transcription factor Sox-2-like, and doublesex- and mab-3-related transcription factor 1-like, were also upregulated with sexual maturation, reflecting active meiotic progression and testicular development. In contrast, genes involved in metabolic processes, including alpha-amylase-like, fatty acid-binding protein brain-like, and fatty acid-binding protein liver-like, were significantly downregulated, indicating shifts in metabolic demands.

In male ganglia, 5-HT1B again showed significant upregulation. This was the only candidate maturation gene to do so. Despite this, several neuropeptide-related genes, such as FMRFamide-like neuropeptide 7 and APGWamide-like neuropeptide, maintained stable and high expression levels. Supplementary Table S7 contains the corresponding expression data for these genes.

### Differentially expressed genes during male gonad development

Again, DESeq2 analysis revealed a large number of genes that were differentially expressed in the gonad during male sexual development. The top 50 differentially expressed genes (adjusted p-value) across male gonadal developmental stages (VGI0 to VGI3) are shown in Figure 5. The expression heatmap separates samples into two distinct clusters, reflecting clear transcriptional differences between immature (VGI0) and more mature (VGI1–VGI3) stages. Most genes exhibit low expression at the immature stage, followed by elevated expression during testis maturation. These patterns imply active transcriptional reprogramming potentially linked to spermatogenesis, structural remodeling, and energy requirements during male gonadal development.

**Figure 5.**
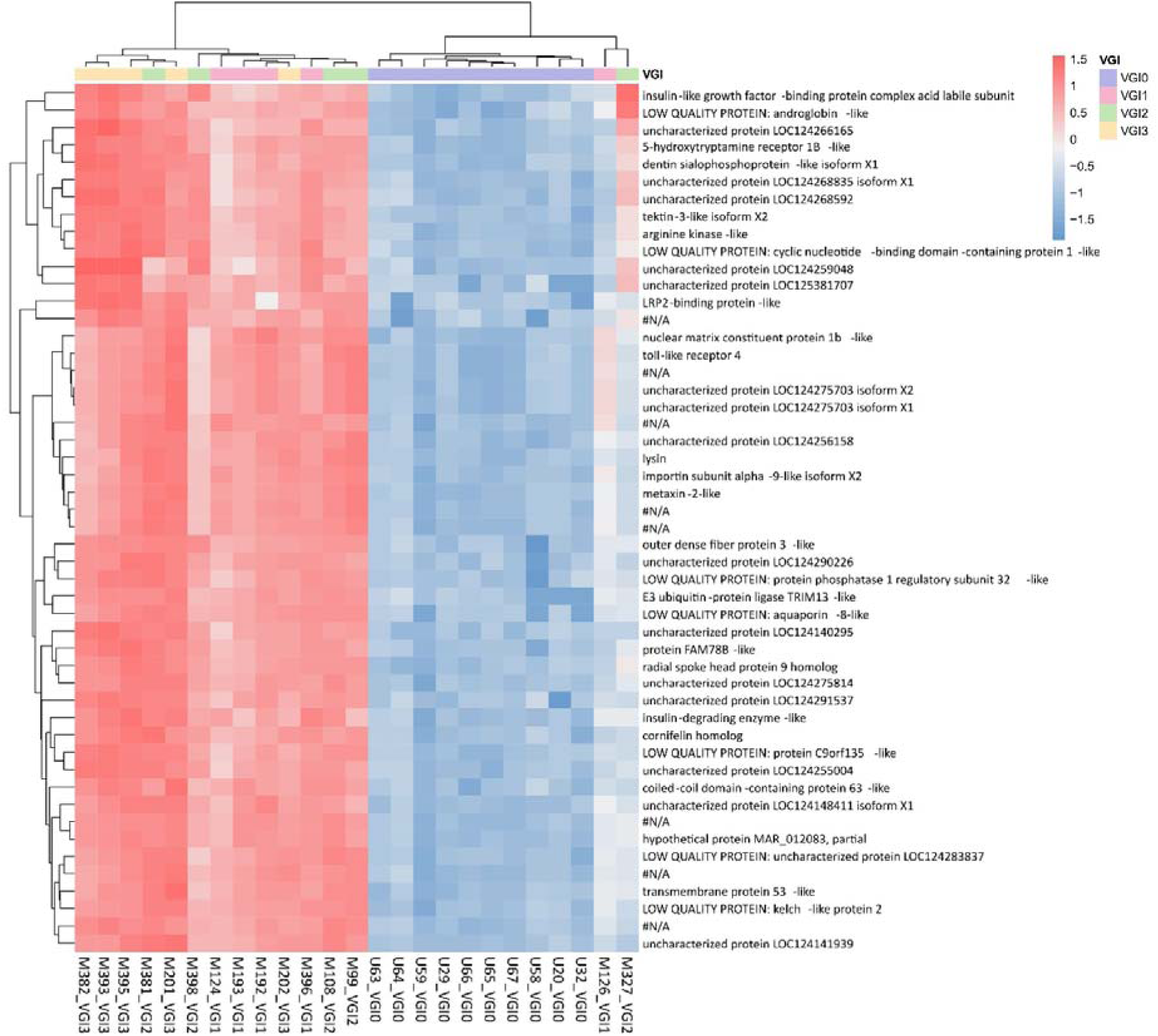
Heatmap showing the expression profiles of the top 50 most differentially expressed genes (adjusted p-value) in male gonads across four developmental stages (VGI0 to VGI3). Samples are hierarchically clustered based on transcript expression similarity. Expression values are color-coded, with blue indicating low expression and red indicating high expression. Developmental stages are color-coded at the top (VGI0: blue, VGI1: pink, VGI2: green, VGI3: orange). The majority of genes display increased expression at VGI1, 2 and 3, suggesting widespread early transcriptional activation during testicular maturation.

All of the top 50 most significantly differentially expressed genes in the male gonad showed increased expression during sexual maturation. Notably, radial spoke head protein 9 homolog and cornifelin homolog were identified among the top 50 DEGs in both male and female gonads, indicating potential shared roles in gonadal development. Several other genes were functionally associated with metabolism, such as arginine kinase-like and insulin-degrading enzyme-like. Genes related to neuroendocrine signaling included 5-HT receptor 1B-like and toll-like receptor 4. Genes implicated in spermatogenesis and meiosis, such as lysin and importin subunit alpha-9-like, exhibited stage-specific expression patterns during testicular development. Additionally, genes that may be involved in immune or stress response pathways were present including TRIM13-like and aquaporin-8-like.

Similar to the pattern observed in female ganglia, several cytochrome P450 family members, particularly those involved in steroid metabolism, exhibited distinct and stage-specific expression during testicular maturation. Notably, cytochrome P450 3A11-like was highly expressed at the pre-maturation stage (VGI0) but significantly downregulated as maturation progressed, suggesting a role in early steroid metabolism or the initiation of gonadal development. In contrast, cytochrome P450 2B1-like showed increased expression at later stages (VGI1–VGI3), indicating a potential function during the mid to late phases of testicular maturation (see Supplementary Table S3).

### Differentially expressed genes in ganglia during male maturation

In order to explore other genes related to sexual maturity in male ganglia, we performed differential expression analysis between developmental stages (VGI1, VGI2, VGI3) which identified additional maturation-associated genes beyond those shown in Figure 6. Notably, several genes were expressed in both the male gonad and male ganglia, showing a significant increase in expression as sexual maturation progressed, including 5-HT receptor 1B-like, toll-like receptor 4, arginine kinase-like, lysin, and importin subunit alpha-9-like isoform X2. Additionally, some genes such as cAMP and cAMP-inhibited cGMP 3’,5’-cyclic phosphodiesterase 10A-like and HORMA domain-containing protein 1-like exhibited increased expression levels during sexual maturation, but were exclusively expressed in male ganglia.

**Figure 6.**
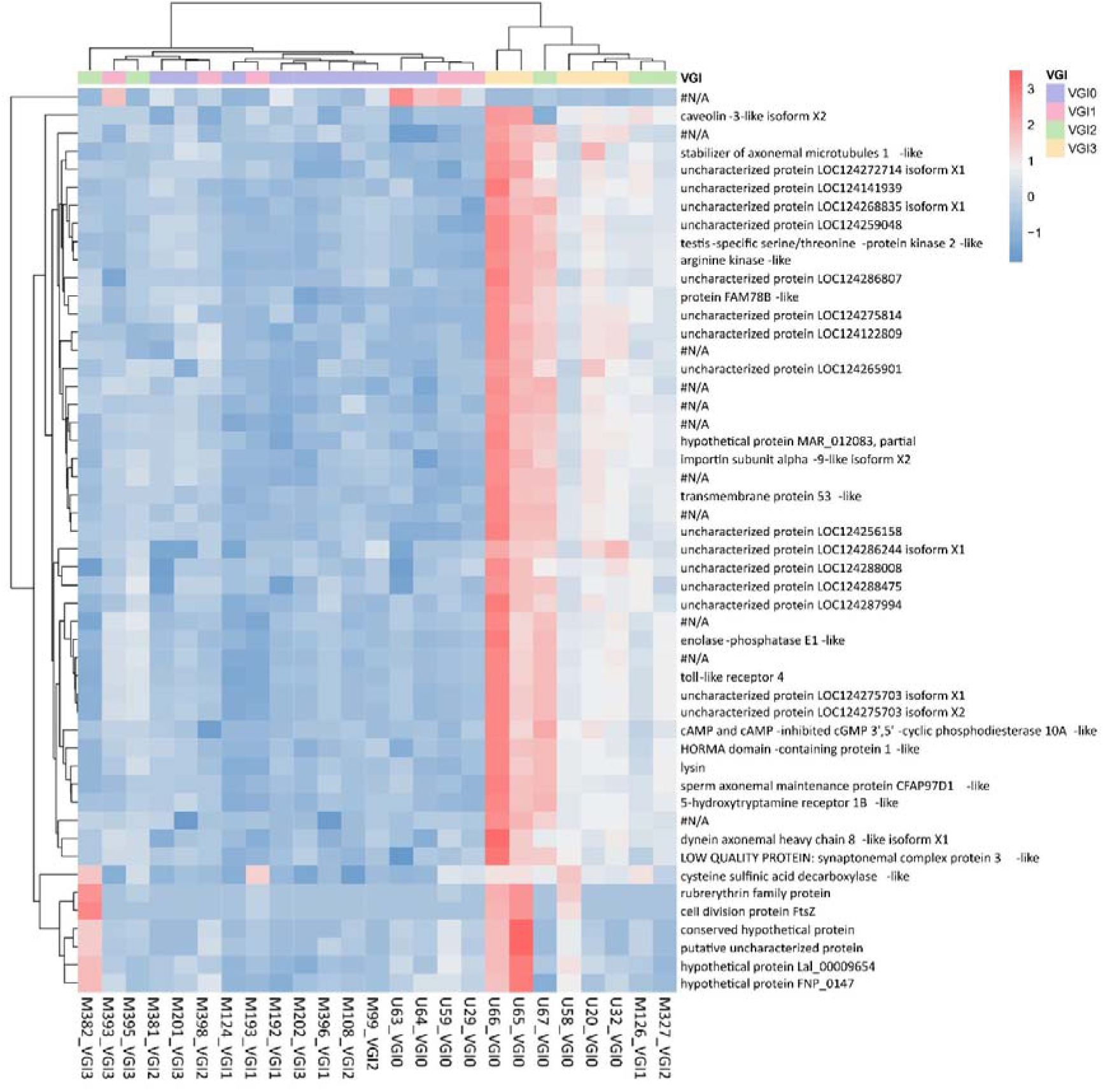
Heatmap showing the expression profiles of the top 50 most differentially expressed genes (adjusted p-value) in male ganglia across four developmental stages (VGI0 to VGI3). Samples are hierarchically clustered based on transcript expression similarity. Expression values are color-coded, with blue indicating low expression and red indicating high expression. Developmental stages are color-coded at the top (VGI0: blue, VGI1: pink, VGI2: green, VGI3: orange). The majority of genes display increased expression at VGI2 and/or VGI3, suggesting stage-specific transcriptional activation in the nervous system during reproductive maturation.

### Gene Network Analysis

To better understand the molecular mechanisms underlying reproductive maturation, we predicted gene regulatory networks using genes that were significantly differentially expressed across maturation stages in female gonad, female ganglia, male gonad, and male ganglia (each separately). From these networks, we extracted key nodes closely associated with sexual maturation, including 5-HT receptor 1B-like, HSP 70 protein and cytochrome P450 3A. Most nodes exhibited one to four connections, indicating a moderately interconnected network. The centrality and interaction patterns of these pivotal genes suggest their participation in core signaling pathways governing maturation.

The subnetworks presented here are functional modules derived from the overall network and emphasize regulatory hubs and pathways relevant to reproductive maturation only. The complete predicted network files are provided as Supplementary File S1-4.

### Predicted Maturation Regulatory Subnetwork in Female Gonad

The 5-HTR1B-like gene was predicted to be a key regulatory gene during maturation in female gonadal tissue (Figure 7) with the subnetwork indicating it potentially regulates 9 downstream genes. This subnetwork was extracted from the gene regulatory network constructed specifically for the female gonad, and includes 5-HTR1B-like, its directly connected nodes (first-degree neighbours), and the nodes connected to those neighbours (second-degree neighbours), representing a local regulatory module. These predicted targets include genes associated with reproductive processes, such as lysin (purple nodes), and those potentially involved in energy metabolism, such as arginine kinase-like (blue-green nodes). A number of subnetwork genes remain uncharacterized (green nodes), suggesting the potential involvement of novel regulatory factors.

**Figure 7.**
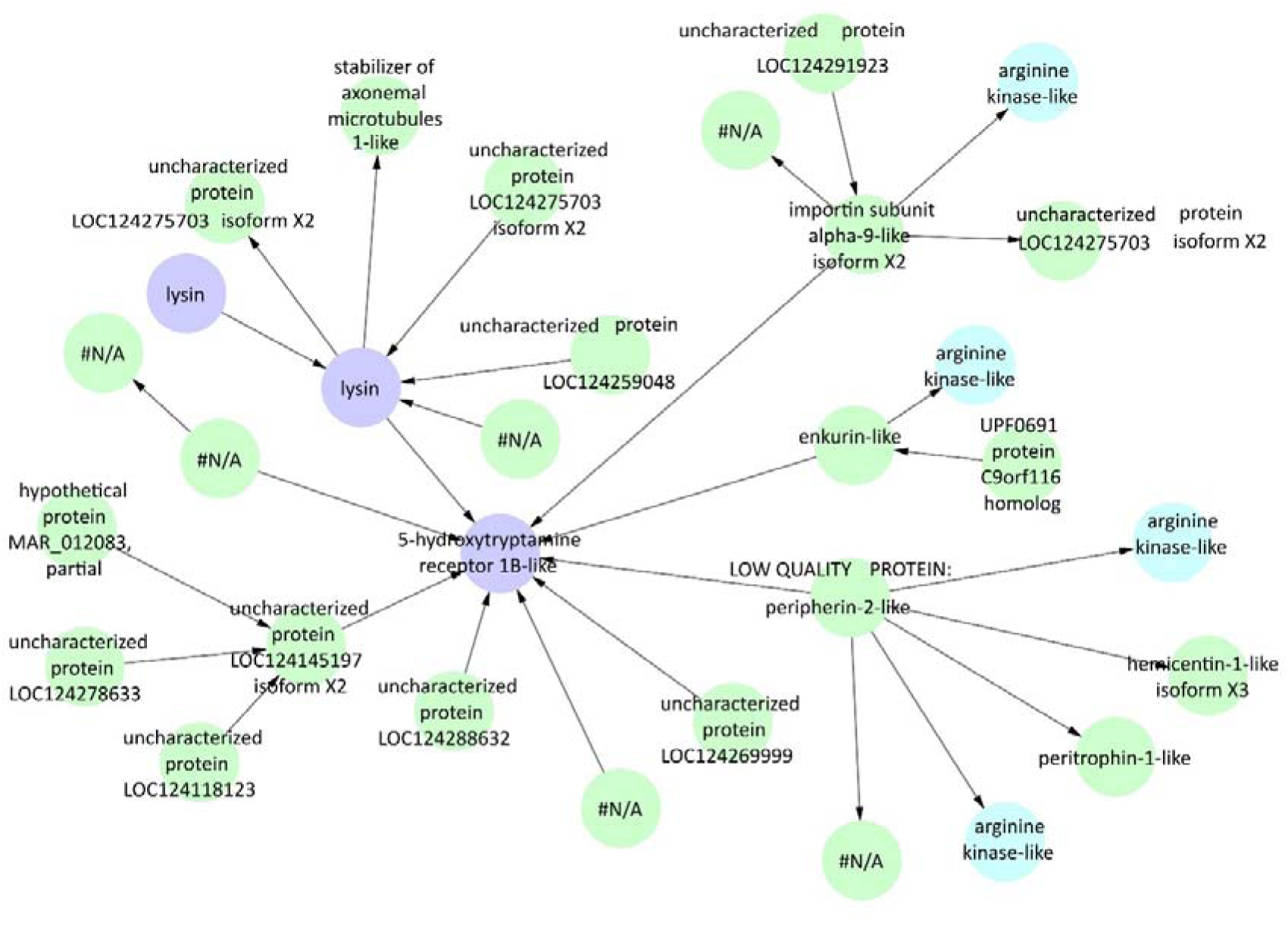
Gene regulatory subnetwork centred on 5-HT receptor 1B-like (5-HTR1B-like) in female gonadal tissue. Purple nodes represent genes associated with sexual maturation, and blue-green nodes represent genes related to energy metabolism. Green nodes indicate other co-expressed genes, including uncharacterized or hypothetical proteins. Arrows (edges) between nodes indicate predicted regulatory relationships.

### Predicted Maturation Regulatory Subnetwork in Male Gonad

5-HT receptor 1B-like was also predicted to be a key regulatory gene during maturation in male gonadal tissue (the subnetwork is illustrated in Figure 8A), with two upstream regulators and one downstream target identified in the predicted regulatory network. This subnetwork was extracted from the gene regulatory network constructed specifically for the male gonad, and encompasses 5-HT receptor 1B-like along with its immediate neighbours and nodes connected by up to three degrees of separation, representing an extended local regulatory module. The predicted interactions highlight its potential role in serotonergic signaling during testis development. The network contains only green nodes beyond the central receptor gene, indicating co-expressed genes not specifically annotated as energy- or maturation-related. These include genes with possible roles in signal transduction (e.g., cAMP and cAMP-inhibited cGMP 3’,5’-cyclic phosphodiesterase 10A-like), cytoskeletal structure (e.g., tubulin alpha-1 chain-like isoform X5, HAUS augmin-like complex subunit 7), apoptosis regulation (O(6)-methylguanine-induced apoptosis 2-like), and cell division. A substantial number of genes remain uncharacterized, highlighting the current gap in functional annotation for marine mollusc genomes. The co-expression of these genes with the 5-HT receptor implies that serotonin signaling may coordinate reproductive, structural, and intracellular signaling processes in the male gonad.

**Figure 8.**
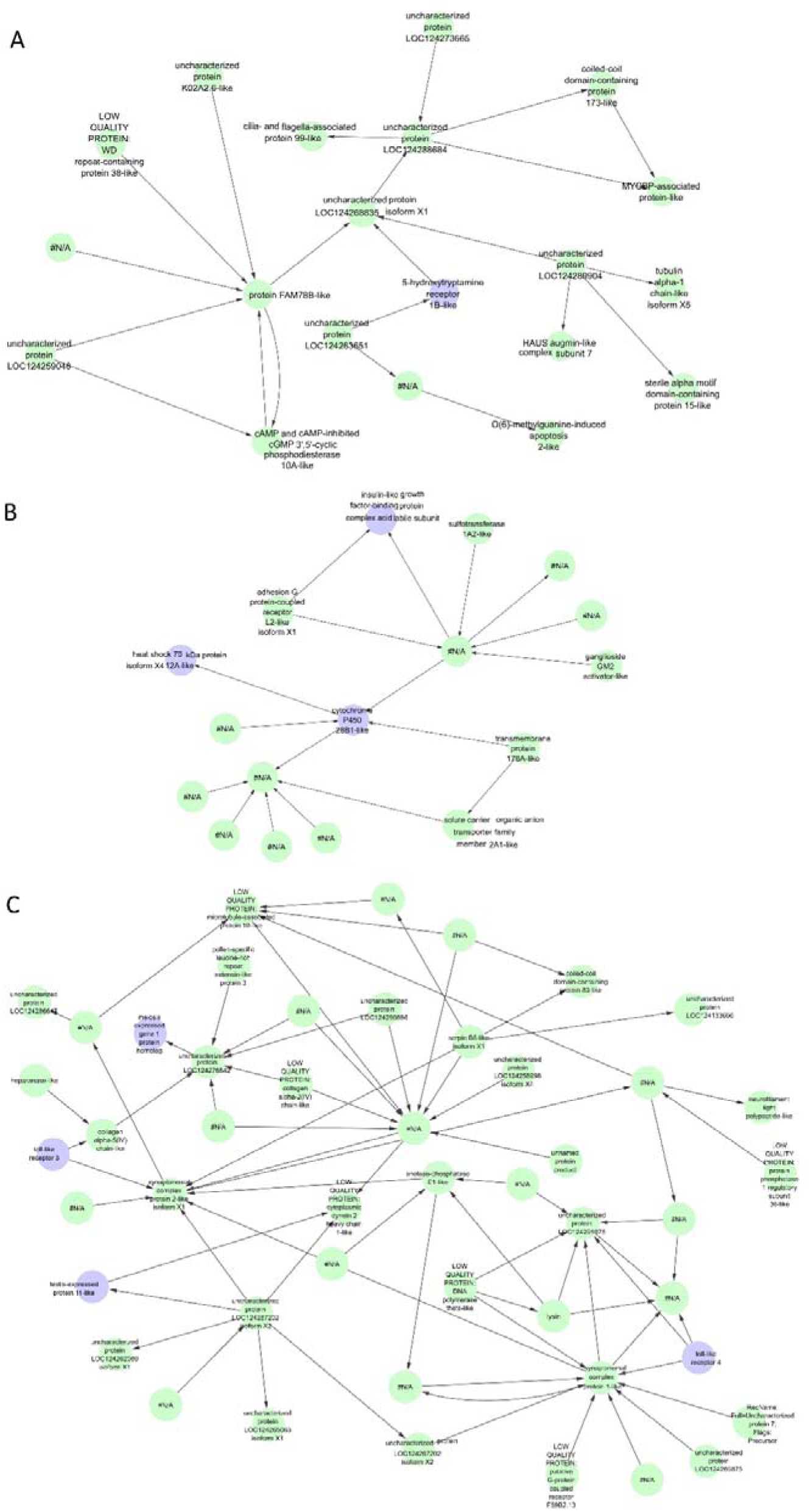
Gene co-expression networks in male gonadal tissue. (a) Network centered on 5-HT receptor 1B-like. (b) Network centered on HSP 70 12A-like isoform X4. (c) Network centered on toll-like receptor 3 and toll-like receptor 4. Green nodes represent co-expressed genes, including both annotated functional genes and uncharacterized proteins. Edges denote putative co-expression relationships inferred from transcriptomic similarity.

HSP 70 12A-like isoform X4 was predicted to play a regulatory role during maturation in male gonadal tissue (the subnetwork is illustrated in Figure 8B). This subnetwork was extracted from the gene regulatory network constructed specifically for the male gonad, and encompasses HSP 70 12A-like isoform X4 along with its immediate neighbours and nodes connected through up to two degrees of separation, representing a local regulatory module potentially involved in stress response or protein homeostasis during testis development. This gene, along with cytochrome P450 26B1-like and insulin-like growth factor-binding protein complex acid labile subunit, has been previously reported to be associated with sexual maturation, and all three are highlighted as purple nodes. The presence of these maturation-related genes suggests a tightly coordinated regulatory module involving stress response, steroid metabolism, and hormonal signaling during testicular development. The remaining co-expressed genes (green nodes) include adhesion G protein-coupled receptor L2-like isoform X1, sulfotransferase 1A2-like, and solute carrier organic anion transporter family member 2A1-like, as well as several uncharacterized proteins. Overall, the network implies that stress-related pathways are closely integrated with reproductive maturation processes in the male gonad.

Figure 8C presents a co-expression subnetwork containing four genes involved in sexual maturation: toll-like receptor 3, toll-like receptor 4, testis-expressed protein 11-like, and meiosis expressed gene 1 protein homolog. These genes are predicted to have linked regulatory functions within the same transcriptional module. These genes are distributed throughout the network but show substantial interconnectivity, suggesting possible transcriptional coordination. Toll-like receptor 3 and 4 imply that immune-related pathways may influence or be influenced by testicular maturation. Testis-expressed protein 11-like and meiosis expressed gene 1 protein homolog highlight the engagement of spermatogenic and meiotic processes. The surrounding green nodes include synaptonemal complex proteins, collagen alpha chain-like genes, and multiple uncharacterized genes, which may contribute to germ cell structural organization or regulatory control. Altogether, this network underscores the integration of immune, structural, and reproductive signaling during male gonadal development.

### Predicted Maturation Regulatory Network in Female ganglia

Figure 9 illustrates the gene regulatory network constructed for the female ganglia, based on 19 genes that were significantly differentially expressed between the VGI0 and VGI2 stages. Edges were filtered in Cytoscape based on interaction strength, retaining the top 40 strongest connections. This resulted in a simplified subnetwork with 17 nodes. Several genes strongly associated with sexual maturation are highlighted in purple, including cytochrome P450 3A24-like, cell division control protein 42 homolog (cdc42), and HSP70 B2-like. These genes are connected to additional genes shown in green, many of which are annotated as uncharacterized or hypothetical proteins. The resulting subnetwork highlights potential regulatory interactions occurring in the ganglia during reproductive development, suggesting that transcriptional changes in the nervous system may contribute to the coordination of female sexual maturation.

**Figure 9.**
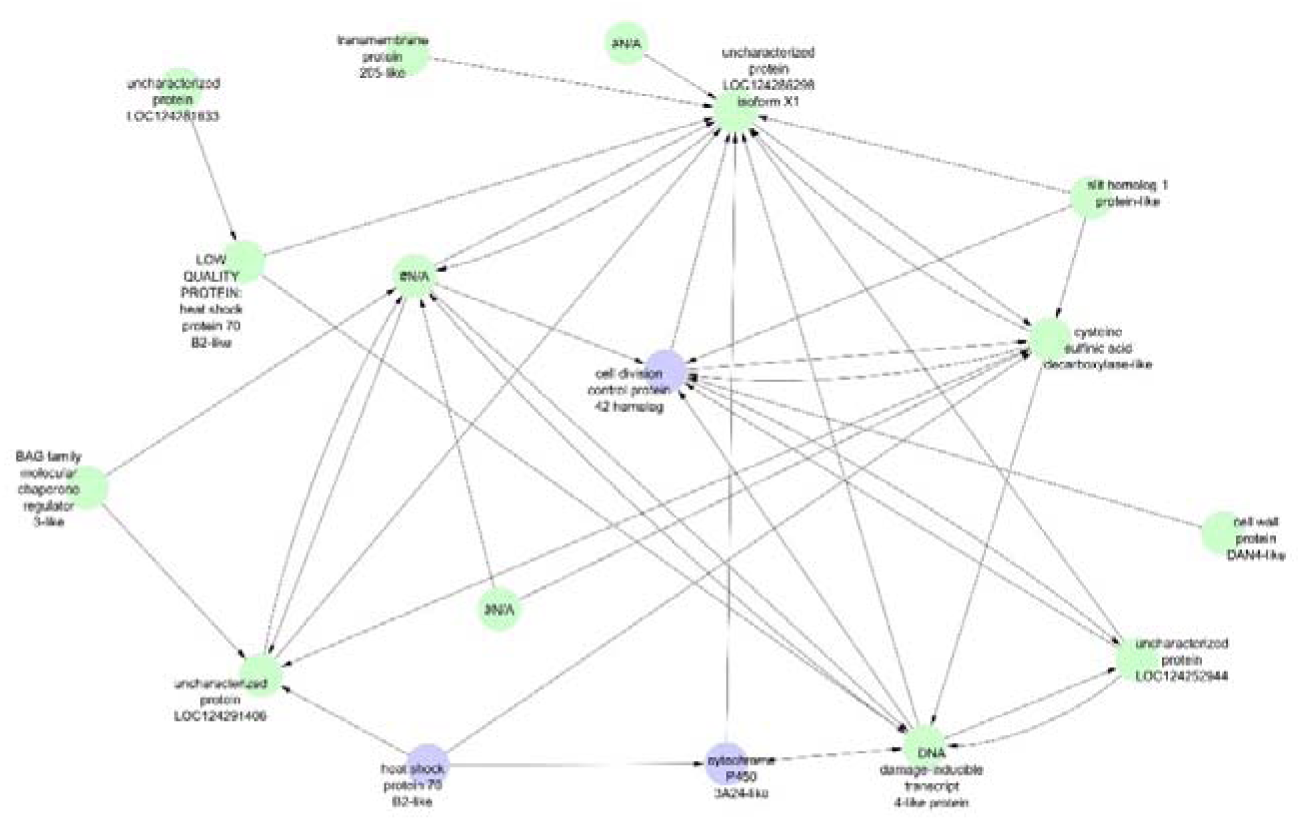
Gene co-expression network of differentially expressed genes in female ganglia between VGI2 and VGI0 stages. Nodes represent genes with significant expression changes; purple nodes indicate genes associated with sexual maturation, and green nodes represent co-expressed genes. Edges indicate putative co-expression or functional associations.

### Predicted Maturation Regulatory Network in Male Ganglia

The 5-HT receptor 1B-like was also predicted to be a key regulatory gene during maturation in male ganglia (the subnetwork is illustrated in Figure 10A), further supporting its conserved regulatory role across tissues. The subnetwork shown was extracted from the gene regulatory network constructed specifically for the male ganglia, and includes 5-HT receptor 1B-like and its directly connected nodes (first-degree neighbours). This gene was again linked to lysin, consistent with its predicted interaction in the female gonadal network. Within this network, three lysin genes (purple nodes) associated with sexual maturation and one arginine kinase-like gene (blue node) involved in energy metabolism were found to be co-expressed with the 5-HT receptor. In addition, numerous green nodes represent other co-expressed genes, many of which are uncharacterized. This network suggests that serotonergic signaling may contribute to sexual maturation by coordinating neural regulation of both reproductive and metabolic pathways in male abalone.

**Figure 10.**
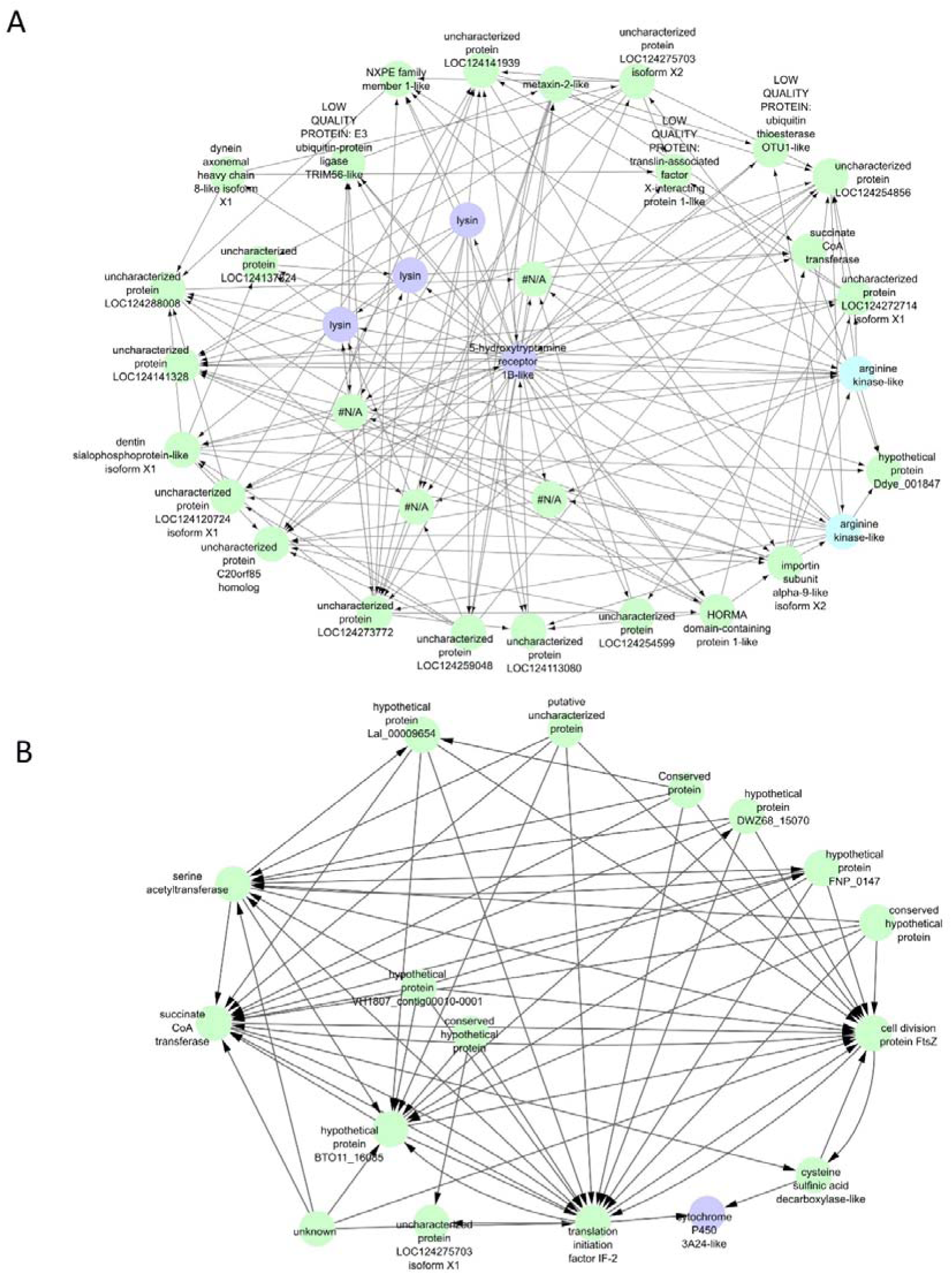
Gene co-expression networks in male ganglia centred on two key genes: (a) 5-HT receptor 1B-like and (b) cytochrome P450 3A24-like. In both networks, edges represent putative co-expression or functional relationships. In panel (a), purple nodes indicate genes associated with sexual maturation, blue nodes represent genes related to energy metabolism, and green nodes denote additional co-expressed genes. In panel (b), the core gene is shown in purple, with green nodes indicating co-expressed genes. These networks suggest that serotonergic signalling and cytochrome P450 pathways may contribute to the coordination of metabolic and reproductive processes in male ganglia.

Cytochrome P450 3A24-like was also predicted to be a regulatory gene during maturation in female ganglia, and was similarly identified as a potential regulator in male ganglia (the subnetwork is illustrated in Figure 10B). This subnetwork was extracted from the gene regulatory network constructed specifically for the male ganglia, and includes cytochrome P450 3A24-like along with its first- and second-degree neighbours. This gene, known for its role in steroid metabolism and reproductive function, was found to be regulated with a range of genes involved in energy metabolism (e.g., succinate CoA transferase, serine acetyltransferase), cell division (cell division protein FtsZ), and translation (translation initiation factor IF-2). The network also includes multiple hypothetical and uncharacterized proteins, suggesting the presence of novel regulatory components. These results imply that cytochrome P450 3A24-like may participate in the neuroendocrine regulation of sexual maturation in male abalone by interacting with genes involved in both reproductive and metabolic pathways.

## Discussion

This study systematically investigated the transcriptional dynamics underlying sexual maturation in *H. laevigata*, revealing a coordinated transition from early-stage energy allocation to later-stage gametogenesis. Differential expression analyses and gene regulatory network prediction uncovered complex interactions among metabolic pathways, hormonal signaling, and reproduction-specific processes across tissues and developmental stages. These findings provide novel insights into the molecular regulation of maturation and suggest that both transient and sustained gene activity are required to orchestrate this multifaceted biological transition.

### Key genes involved in sexual maturation

#### 5-hydroxytryptamine receptor 1B-like

5-HT also known as serotonin, is a key neurotransmitter that plays an essential role in the neuroendocrine regulation of reproduction across invertebrates. It promotes oocyte maturation, gamete release, and gonadal development via its receptors (Osada et al., 1998, Zatylny et al., 2000, Stricker and Smythe, 2001). In molluscs and crustaceans 5-HT stimulates the release of gonad-regulating hormones and activates androgenic glands in males (Fingerman, 1997, Sharker et al., 2020). In bivalves, 5-HT has been shown to modulate gamete maturation through serotonergic neurons projecting to gonadal tissues, with receptor expression upregulated by 17β-estradiol (Fingerman, 1997, Sharker et al., 2020, Panasophonkul et al., 2009). In abalone specifically (*H. discus hannai*, *H. asinina*), 5-HT receptors are widely expressed in ganglia and increase during ovarian maturation (Wang and He, 2014). In this study, *H. laevigata* exhibited elevated 5-HT receptor expression during maturation in female and male gonads, and in male ganglia, while no significant increase was observed in female ganglia. These results suggest a conserved but sexually distinct role for 5-HT in molluscan reproductive regulation (Wang and He, 2014).

In the gene network analysis, 5-HT receptor 1B-like was identified as a key regulatory node in both male and female gonads, as well as in male ganglia, and was predicted to regulate key downstream genes such as lysin and arginine kinase-like. Although no significant differential expression was detected in female ganglia, this gene remained stable and highly expressed throughout development, suggesting a potential constitutive role. Both lysin and arginine kinase-like exhibited markedly increased expression following the onset of sexual maturation in male and female gonads, highlighting a conserved regulatory role for 5-HT receptor 1B-like in coordinating reproduction-related gene expression across tissues.

Lysin is a non-enzymatic protein specifically expressed in the acrosome of abalone sperm, where it binds to the vitelline envelope (VE) receptor on the egg surface to facilitate sperm entry and fertilization (Swanson and Vacquier, 1997, Vacquier and Lee, 1993). It exhibits species-specific activity, primarily determined by sequence variability at the N-terminus (Shaw et al., 1994), and has undergone strong positive selection across abalone species, likely driven by sperm competition (Swanson et al., 2001). While traditionally studied in the context of male fertilization, this study found that lysin genes were significantly upregulated in both male and female gonads during sexual maturation, with notably higher expression in males (see Supplementary Table 1 and 3). Similar dual-sex expression has been reported for lysin-like proteins in mussels (*Mytilus edulis*), where acrosomal proteins such as M6 and M7 are also expressed in female gametes and somatic tissues. These findings suggest that lysin may contribute not only to fertilization but also to broader roles in gamete maturation and reproductive readiness (Heß et al., 2012).

Building on the gene network predictions, 5-HT receptor 1B-like was inferred to modulate the expression of multiple lysin genes, implying that serotonergic signalling may play a broader role in reproductive coordination. Given the classical function of lysin in sperm acrosome reactions, its notable expression in ganglia, together with its predicted regulation by 5-HT receptor 1B-like, suggests it is involved in the neuroendocrine control of gamete release, sperm maturation, or reproductive behaviour. This expands the known scope of lysin beyond fertilization and underscores the neural-reproductive interface in abalone maturation. Further experimental validation, such as transcription factor binding assays and in situ protein localization, is needed to confirm the functional relevance of this interaction.

#### Cytochrome P450 family

The cytochrome P450 (CYP) superfamily is involved in a range of metabolic pathways, including fatty acid metabolism and detoxification (Van Bogaert et al., 2011, Lu et al., 2021). In the context of reproductive development, its role in steroid metabolism is particularly relevant, with CYP19A1 (aromatase) being essential for the conversion of androgens into estrogens in vertebrates (Niwa et al., 2015, Auchus and Miller, 2015). Although aromatase-like activity and invertebrate-type steroids have been reported in molluscs (Hallmann et al., 2019, Scott, 2012), other studies suggest that the detected vertebrate-type steroids may result from environmental uptake rather than endogenous synthesis (Schwarz et al., 2017). These findings imply that other CYP subtypes may carry out analogous or mollusc-specific steroid-modifying functions that contribute to reproductive maturation in molluscs (Niwa et al., 2020).

In the male gonads of *H. laevigata*, CYP3A11-like was highly expressed at the pre-maturation stage (VGI0) and significantly downregulated as sexual maturation progressed. This expression pattern is consistent with observations in sea urchins (*Mesocentrotus nudus*), where CYP3A homologs have also been implicated in early-stage gonadal development and show decreasing expression as gonads mature (Su et al., 2024). These findings suggest that CYP3A11-like may act as a molecular switch involved in the initial regulation of steroid biosynthesis and maturation onset in abalone.

In contrast, no significant differential expression of CYP3A genes was observed in the female gonads (Figure 1). However, cytochrome P450 12A2-like and cytochrome P450 4F6-like showed relatively high expression levels prior to maturation, followed by reduced expression after maturation was initiated, although these changes were not statistically significant. This sex-specific expression pattern may indicate distinct steroid regulatory pathways between males and females, possibly reflecting differential timing or hormonal sensitivity during reproductive development.

In both female and male ganglia, CYP3A expression exhibited a gradual increase from immature to mature stages, suggesting a potential role in the neuroendocrine regulation of reproduction. While the involvement of CYP genes in gonadal steroidogenesis is well established, their expression and function in nervous tissues remain largely unexplored in molluscs. Given the observed patterns, it is plausible that CYP3A participates in steroid hormone metabolism within ganglia, thereby indirectly influencing gonadal development through neuroendocrine signaling.

Another CYP gene, cytochrome P450 2B1-like, displayed an opposite trend: it was lowly expressed at early stages and significantly upregulated in male gonad after maturation began (VGI1 to VGI3). Similar patterns of stage-specific CYP3A expression have been reported in mussels (*M. edulis*) during reproductive development (Cubero-Leon et al., 2012, Giuliani et al., 2013), implying that different CYP family members may function sequentially throughout maturation.

Gene regulatory network prediction revealed a potential regulatory interaction in male gonads, in which HSP70 12A-like isoform X4 is predicted to regulate the expression of cytochrome P450 2B1-like. Although interactions between HSPs and CYP450s have previously been reported in the context of stress responses and xenobiotic metabolism (Snyder et al., 2001), the presence of this link in a reproductive maturation network raises the possibility that it may also play a role in gonadal development. This finding highlights a potential intersection between stress-related pathways and reproductive signaling in abalone.

#### Heat shock protein 70

The HSP70 family plays a critical role in reproductive development by responding to cellular stress and maintaining protein homeostasis. In invertebrates, HSP70 shows stage-specific patterns. In *Octopus tankahkeei* and *Cherax quadricarinatus* it localizes specifically to germ cells during spermatogenesis but is absent in mature sperm and somatic cells (Long et al., 2015, Fang et al., 2012). In *Magallana gigas* HSP70 expression is elevated during early spermatogenesis but decreases in spermatozoa, while remaining high in female oocytes (Kingtong et al., 2013).

In this study, HSP70 12A-like was progressively upregulated in male greenlip abalone during sexual maturation. In females, HSP70 12A-like was absent, while HSP70 4-like remained stably expressed across all stages. These findings suggest isoform-specific roles of HSP70 in early germ cell development in *H. laevigata*, warranting further research into its regulatory mechanisms during reproduction in marine invertebrates.

#### Toll-like receptors

Toll-like receptors (TLRs), as key members of the pattern recognition receptor (PRR) family, are primarily known for their role in recognizing pathogen-associated molecular patterns (PAMPs) and initiating innate immune responses (Wang et al., 2011). While their involvement in immune defense is well established, their role in reproductive development, particularly in invertebrates, remains largely unexplored. In vertebrates, however, TLRs have emerged as important regulators of reproductive processes (Girling and Hedger, 2007). Current research on TLRs in molluscs is primarily focused on their roles in immune responses (Saco et al., 2023, Xu et al., 2019). In females, TLRs contribute to ovarian, endometrial, and placental functions and have been associated with reproductive complications such as intrauterine growth restriction and preterm birth (Girling and Hedger, 2007). In males, TLRs have been linked to the regulation of testicular steroidogenesis and spermatogenesis, suggesting broader roles in male reproductive physiology (Girling and Hedger, 2007).

In this study, we found that TLR3 was highly expressed in the male gonads of *H. laevigata* prior to sexual maturation (VGI0), but its expression declined after maturation commenced. In contrast, TLR4 expression increased during the post-maturation stages. Although similar expression trends were observed in female gonads, the differences were not statistically significant. Notably, gene network analysis revealed that both TLR3 and TLR4 were associated with meiosis expressed gene 1 protein homolog and testis-expressed protein 11-like in male gonads, indicating that TLR signaling may be involved in the regulation of spermatogenesis and testicular development. These findings suggest a potential reproductive role for TLRs beyond their canonical immune functions in greenlip abalone.

#### Gonadotropin-releasing hormone receptor, GnRH

Gonadotropin-releasing hormone (GnRH) is a key hypothalamic neurohormone that regulates reproductive function in both vertebrates (Hazum and Conn, 1988, Chen and Fernald, 2008) and invertebrates (Rastogi et al., 2002). Its precursor undergoes enzymatic cleavage and post-translational modifications to generate the mature hormone (Harris, 1989). Although GnRH-like peptides have been identified in various invertebrates, their structure and function may differ significantly due to evolutionary divergence (Sakai et al., 2017). In molluscs, GnRH or GnRH-like peptides have been reported in species such as mussels (*M. edulis*) (Mathieu et al., 1988), scallops (*Patinopecten yessoensiss*) (Nakamura et al., 2007), Pacific oyster (*C. gigas*) (Bigot et al., 2012), and Pacific abalone (*H. discus hannai*) (Funayama et al., 2019), with strong evidence supporting their regulatory roles in gonadal development.

In this study, GnRH-like peptide showed significantly increased expression in female gonads during early sexual maturation stages (VGI1 and VGI2), consistent with findings in *Ruditapes philippinarum*, where expression peaks at the onset of sexual maturity (Song et al., 2015). However, it was not identified as a potential regulatory node in the corresponding gene network analysis. This absence is likely attributable to the construction of the network based solely on differentially expressed genes; potential interacting partners of GnRH-like that did not meet the threshold for differential expression were excluded, thereby precluding the inclusion of GnRH-like in the inferred network topology. In contrast, male gonads displayed stable GnRH-like peptide expression throughout all maturation stages (VGI0–VGI3), suggesting a more constant or baseline regulatory function in males. These results imply that GnRH may play sex-specific roles in abalone reproduction. Future studies are needed to elucidate the mechanisms underlying GnRH function in molluscan sexual maturation and to further explore its potential as a target for reproductive regulation.

#### Energy storage in early maturation

Energy metabolism plays a crucial role in supporting early gonadal development, with specific genes involved in carbohydrate and lipid processing showing dynamic expression patterns during sexual maturation. The α-amylase catalyzes the hydrolysis of α-1,4-glycosidic bonds in starch to produce glucose, which is subsequently converted to glycogen and stored for use during reproductive development (Henrissat, 1991, Kasar et al., 2022, Guzmán-Maldonado et al., 1995, Andreotti et al., 2006, Satoh et al., 2013). In this study, α-amylase-like was highly expressed in both male and female gonads at the early stage of sexual maturation (VGI0), with a marked decline observed in later stages (VGI2/VGI3), suggesting its involvement in energy accumulation prior to gametogenesis. Similar trends have been observed in *M. gigas* and *Argopecten irradians*, where energy reserves are established before vitellogenesis (Li et al., 2024, Barber and Blake, 1981). In *M. gigas*, RNA interference targeting *α-amylase* led to reduced gonad size and germ cell proliferation (Huvet et al., 2015). However, a contrasting pattern was reported in *H. discus hannai*, where amylase expression peaks at mature stages (Kim et al., 2020) , indicating species- or stage-specific variation in regulation.

Fatty acid-binding proteins (FABPs) are cytosolic carriers of hydrophobic molecules such as fatty acids and lipids, playing essential roles in lipid transport and metabolism (Storch and Thumser, 2000). In this study, three FABP isoforms (brain-like, liver-like, and an uncharacterized variant) were strongly expressed at VGI0 but downregulated as maturation progressed, suggesting early involvement in lipid accumulation. Similar early-stage FABP expression has been documented in *Solea senegalensis* and scallops, where lipids serve as critical energy sources for gonadal development (McKeegan and Sturmey, 2011, Agulleiro et al., 2007). In contrast, late-stage FABP upregulation has been reported in *Scylla paramamosain* and *H. discus hannai* (Zeng et al., 2013, Kim et al., 2020). Additionally, yolk ferritin-like, an iron-storage gene, was co-expressed with *FABP-brain-like* in the gene network and showed a parallel expression trend, implying a cooperative role in coordinating iron and lipid reserves for oocyte development (Bottke, 1982, Brooks and Wessel, 2002). These findings underscore the critical function of α-amylase-like and FABP genes in energy provisioning during the initial stages of sexual maturation.

#### Early maturation regulatory genes

This study identified several genes potentially involved in the regulation of early sexual maturation in greenlip abalone (*H. laevigata*). Among these, 5-HTR1B-like emerged as a key regulatory node, showing upregulation after maturation began in the female gonad, male gonad, and male ganglia, while maintaining stable expression in female ganglia. Its predicted downstream targets, including lysin and arginine kinase-like, were significantly upregulated during maturation, suggesting a broader role for serotonergic signaling in coordinating gamete development and reproductive readiness across neural and gonadal tissues.

Cytochrome P450 genes exhibited distinct sex- and tissue-specific expression patterns. CYP3A11-like was highly expressed in immature male gonads and downregulated during maturation, indicating a potential early role in steroid metabolism. In contrast, CYP2B1-like was upregulated at later stages in the male gonad. The increased expression of CYP genes in ganglia further suggests possible neuroendocrine roles beyond their classical functions in gonadal tissues, indicating that different CYP family members may act at different stages of sexual maturation.

Together, these findings highlight the coordinated involvement of neuroendocrine, metabolic, and immune-related genes in the early stages of abalone sexual maturation, and provide a foundation for future functional and comparative studies in molluscan reproductive biology.

While the results offer valuable advances in our understanding of maturation, several factors should be considered when interpreting these findings. The absence of late-stage (VGI3) female samples may limit the completeness of stage-specific comparisons between sexes. In addition, many gene functions were inferred through computational annotation pipelines, which are subject to inaccuracies, particularly in non-model species like abalone. A substantial proportion of differentially expressed genes also remain uncharacterized, constraining biological interpretation in some cases. Moreover, although gene network analyses provide useful predictions of regulatory relationships, they do not establish causality and should be interpreted with caution. These caveats highlight the need for further validation and expanded datasets in future studies.

## Conclusion

This study provides novel insights into the transcriptional regulation of sexual maturation in *H. laevigata*, particularly highlighting the early activation of genes involved in germ cell proliferation, meiotic initiation, and neuroendocrine signaling. These early-expressed genes appear to play foundation roles in initiating reproductive maturation and establishing a regulatory framework for subsequent developmental processes. While early-stage gonads also exhibited elevated expression of genes linked to energy metabolism, these likely serve as supportive mechanisms to sustain the biosynthetic demands of reproductive activation. Together, these findings underscore the importance of early-stage transcriptional programs in directing sexual maturation and offer a valuable molecular resource for further functional studies and aquaculture applications. Given the conserved nature of many of the identified pathways, these insights are likely to be relevant to other *Haliotis* species and potentially to molluscan reproduction more broadly.

## Supplementary Data

Supplementary data to this article can be found online at https://doi.org/10.25919/d9f3-0435.

## Ethical statement for experimental animals

All animal tissues used in this study were repurposed from the Australia Government Seafood CRC Grant 2010/767.

## CRediT authorship contribution statement

Ya Zhang - Designed the experiments, conducted the experimentation, performed data analysis, and wrote the manuscript; Carmel McDougall - Designed the experiments, performed data analysis, and reviewed the manuscript; Ido Bar - Performed data analysis, and reviewed the manuscript; Natasha Botwright - Collected samples, designed the experiments, performed data analysis, and reviewed the manuscript.

## Acknowledgements

We thank David Connell (Kangaroo Island Abalone) for supplying the abalone from the original study in 2010 to 2011, and Yumbah Aquaculture for their support of this project.

## Data availability

The raw sequencing data generated and analyzed during this study have been deposited in the NCBI Sequence Read Archive (SRA) under BioProject accession number [PRJNA1242192]. The transcriptome assembly, annotation, transcriptome analysis statistics, and the R code used for differential gene expression analysis and figure preparation are available for download from the CSIRO Data Access Portal (Zhang et al., 2025).

