## Supplementary Table S1 for "Characterization of the molecular mechanisms of early sexual maturation stages in the Australian greenlip abalone (*Haliotis laevigata*)"

Supplementary Table S1. Candidate genes for sexual maturation with associated species and tissues.

| Gene Name | ***Species*** | Sex | | Tissue | | Reference |
|---|---|---|---|---|---|---|
| | | Female | Male | Gonad | Ganglia | |
| 17-beta-hydroxysteroid dehydrogenase | *Mizuhopecten yessoensis* | * | * | * | | [1] |
| 3 beta-hydroxysteroid dehydrogenases | *M. yessoensis* | * | * | * | | [1] |
| 5-hydroxytryptamine receptor | *Haliotis discus hannai* | * | | * | | [2] |
| achatin-like | *H. discus hannai* | * | | | * | [3] |
| alpha-amylase | *H. discus hannai* | * | | * | | [4] |
| APGWamide-like neuropeptide | *H. discus hannai* | * | | | * | [5] |
| | *H. asinina* | | * | * | | [6] |
| buccalin-like | *Saccostrea glomerata* | * | * | | * | [7] |
| doublesex- and mab-3-related transcription factor | *Tridacna crocea* | | * | * | | [8] |
| Crustacean Cardioactive Peptide | *H. discus hannai* | * | | | * | [3] |
| cytochrome P450 family | *Mesocentrotus nudus* | * | * | * | | [9] |
| egg laying hormone | *H. laevigata* | * | | * | | [10] |
| estradiol 17-beta-dehydrogenase | *H. diversicolor supertexta* | * | | * | | [11] |
| fatty acid-binding protein | *H. discus hannai* | * | | * | | [4] |
| FMRFamide-like neuropeptide | *H. discus hannai* | * | | | * | [3] |
| forkhead box protein | *Magallana (Crassostrea) gigas* | * | | * | | [12] |
| Gonadotropin-releasing hormone | *H. asinina and H. laevigata* | * | | | * | [13] |
| heat shock 70 kDa protein | *H. laevigata* | * | | * | | [14] |
| insulin-like growth factor | *H. asinina* | * | * | | * | [15] |
| meiosis related genes | *M. gigas* | * | * | * | | [12] |
| myomodulin neuropeptide 1-like | *H. asinina* | * | * | | * | [15] |
| pedal peptide A | *H. discus hannai* | * | | | * | [3] |
| pedal peptide B | *H. discus hannai* | * | | | * | [3] |
| Sperm related proteins | *M. gigas* | | * | * | | [16] |
| transcription factor Sox | *H. discus hannai* | * | * | * | | [17] |
| vasa-like protein | *H. discus hannai* | * | * | * | | [18] |
| vitelline envelope sperm lysin receptor | *H. rufescens* | * | | * | | [19] |
| whitnin | *H. asinina* | * | * | | * | [15] |

* Indicates that the gene was reported to be associated with the corresponding sex or tissue type in the referenced study.
