## Supplementary figures and images for "Characterization of the molecular mechanisms of early sexual maturation stages in the Australian greenlip abalone (*Haliotis laevigata*)"

### Supplementary Figure S1

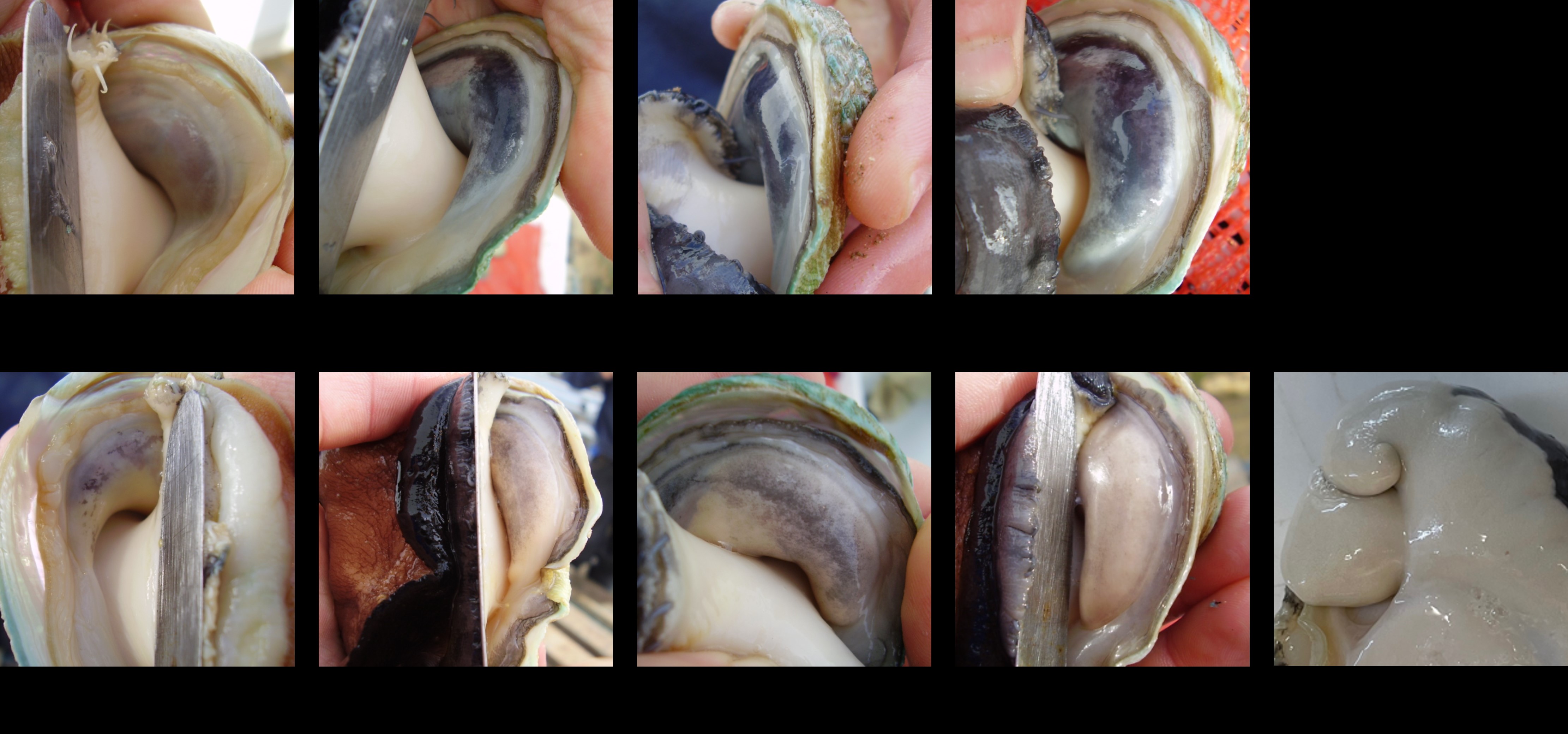
